# Characterizing Surround Suppression with Dynamic Natural Scenes

**DOI:** 10.1101/2025.08.11.669612

**Authors:** Merve Kiniklioglu, Daniel Kaiser

## Abstract

Surround suppression refers to the reduction in perceptual sensitivity to a central stimulus caused by its surrounding context. Previous studies have examined this phenomenon mainly with simple stimuli varying only in low-level visual features, leaving it unclear whether the same principles apply to natural scenes. To address this, we investigated surround suppression in complex and dynamic scenes by systematically manipulating categorical similarity and motion congruence between the center and surrounding scenes. Categorical similarity was examined across four levels: identical exemplars, different exemplars of the same basic-level category, different basic-level categories, and different superordinate categories. Motion direction was manipulated by center and surround drifting either in the same or opposite directions. In two experiments, we measured contrast sensitivity for the center scene during a categorization task. We found that the suppression increased as the categorical similarity between the center and the surround decreased, with the strongest suppression observed for different superordinate categories. This contrasts with results for simple stimuli, where increasing center-surround similarity increases suppression. Yet, consistent with the findings from simple stimuli, suppression was stronger when the center and surround moved in the same direction. These findings show that contextual modulation in natural vision is governed not only by low-level feature similarity but also by high-level categorical structure. Context-dependent suppression may therefore help the visual system prioritize coherent, task-relevant information while filtering incongruent input.

## Introduction

Visual perception is inherently context-dependent. The appearance and detectability of a stimulus can be significantly influenced by surrounding visual information (Tadin et al., 2003). One well-known example of such contextual modulation is surround suppression, in which the presence of surrounding stimuli reduces perceptual and neural sensitivity to a central target (Cavanaugh et al., 2002a, 2002b; DeAngelis et al., 1994; Ichida et al., 2007; Petrov et al., 2005; Pihlaja et al., 2008; Shushruth et al., 2012, 2013; Walker et al., 1999; Williams et al., 2003; Xing & Heeger, 2001; Zenger-Landolt & Heeger, 2003). Surround suppression has been extensively studied using simple stimuli, such as static or drifting gratings, and has been shown to depend on low-level features, including contrast, orientation, and motion direction (Angelucci et al., 2017; Cavanaugh et al., 2002b; Er et al., 2020; Kiniklioglu & Boyaci, 2022; Schallmo et al., 2018; Tadin, 2015; Turkozer et al., 2016). Suppression is strongest when the center and surround share similar features, but it weakens or even reverses to facilitation when they differ. For example, iso-oriented surrounds typically suppress responses to a central stimulus, whereas orthogonal surrounds can lead to facilitation (Cavanaugh et al., 2002a; DeAngelis et al., 1994; Flevaris & Murray, 2015; Schallmo et al., 2016; Serrano-Pedraza et al., 2012; Shushruth et al., 2012; Walker et al., 1999). A similar pattern is observed with motion direction: when center and surround gratings drift in opposite directions, suppression is reduced compared to when they move in the same direction (Allman et al., 1985; Born & Tootell, 1992; Cavanaugh et al., 2002b; Kastner et al., 1995; Kiniklioglu & Boyaci, 2025; Kiniklioglu & Boyaci, 2022; Lamme, 1995; Paffen, Alais, & Verstraten, 2005; Paffen, van der Smagt, et al., 2005). Although the characteristics of surround suppression for simple stimuli are well established, it remains unclear how surround suppression influences the processing of more complex naturalistic stimuli. Animal studies using more ecologically relevant stimuli have yielded conflicting results. Coen-Cagli et al. (2015) reported stronger surround suppression in the macaque primary visual cortex when the center and surround consisted of homogeneous natural images. In contrast, Onat et al. (2013) found that in natural scene videos, surround patterns that perceptually complete the central stimulus elicit facilitation in the cat’s primary visual cortex. Extending these findings, Fu et al. (2024) used convolutional neural networks (CNNs) trained on neural recordings from mouse visual cortex to model center-surround interactions for natural scenes. Their results showed that congruent center-surround pairings yielded excitatory effects, whereas incongruent pairings yielded inhibitory effects. Taken together, the findings obtained in animals paint a complex and somewhat inconclusive picture of center-surround interactions in natural vision. Whether center-surround similarity leads to suppression or facilitation in naturalistic contexts is still not fully understood. While surround suppression with natural stimuli has not been systematically investigated in humans, high-level contextual influences on perception have been studied. In particular, the categorical relationship between central and peripheral content, such as whether both belong to the same scene category (e.g., indoor vs. outdoor), has been shown to influence perceptual judgments in scene categorization tasks (Faurite et al., 2024; Lukavský, 2019; Trouilloud et al., 2020; Trouilloud et al., 2022). For example, categorization accuracy declines when the surrounding context belongs to a different category, even when it is unattended (Lukavský, 2019). Such categorical congruency effects interact with physical stimulus characteristics: Peripheral low spatial frequency information facilitates central scene categorization when it is categorically congruent (Faurite et al., 2024). In contrast, incongruent peripheral scenes impair performance, especially when central and peripheral scenes share similar physical properties, such as amplitude spectrum and spatial configuration. Notably, even categorically congruent surrounds can impair central scene categorization when they are physically dissimilar from the center (Peyrin et al., 2021). Taken together, studies on scene congruency effects suggest an interplay of high-level characteristics and low-level visual attributes. This raises an important question: Does surround suppression for natural stimuli follow the same principles observed with low-level visual stimuli, or is it governed by fundamentally different mechanisms, potentially driven by high-level stimulus characteristics?

To address this question, we investigated whether surround suppression in dynamic natural scenes is modulated by high-level categorical context. To do so, we measured contrast sensitivity to a central scene while systematically manipulating the categorical similarity between the center and surrounding scenes across four levels: (1) identical exemplar, (2) different exemplar from the same basic-level category, (3) different basic-level categories within the same superordinate category, and (4) different superordinate categories. This design allowed us to assess how increasing categorical distance influences the perceptual strength of suppression. Additionally, we manipulated the motion congruency between the center and surround, specifically whether they drifted in the same or opposite directions, to test whether established low-level motion effects observed with simple stimuli generalize to more naturalistic stimuli. By jointly manipulating categorical similarity and motion congruence, our study characterizes how high- and low-level contextual factors shape surround suppression in dynamic natural scenes.

## Experiment 1

In Experiment 1, we investigated whether categorical similarity between a central scene and its surrounding context influences contrast sensitivity for the central scene. Participants performed a scene categorization task in which the contrast of both the central and surrounding scenes was adjusted adaptively. This allowed us to estimate contrast sensitivity under varying contextual conditions, comparing performance across four levels of categorical similarity (identical exemplar, different exemplar, different basic-level category, different superordinate category) and two motion congruency conditions ( same-direction, opposite-direction).

## Methods

### Participants

Twenty-two volunteers (11 male, mean age= 27.6 years), participated in Experiment 1. All participants reported normal or corrected-to-normal vision. They gave written informed consent prior to the experiment and received monetary compensation for their participation. The study was approved by the Ethics Committee of Justus Liebig University Giessen. All experimental protocols were in accordance with the Declaration of Helsinki.

### Stimuli

The stimuli consisted of panoramic videos created by moving static scene images behind a circular occluder (520 × 400 pixels, 3 pixels per frame). The center images were displayed through a 1.9°-diameter aperture, while the surrounding images were presented through an annular aperture ranging from 2.5° to 10.4° in diameter. To separate the center and surrounding scenes, the area between 1.9° and 2.5° remained unstimulated, creating a gap between the center and annular stimuli. To ensure smooth transitions and minimize sharp contrast differences at the aperture edges, a cosine envelope was applied to smooth the boundaries between the center and surround, as well as between the outermost edge of the surround and the background. All images were converted to grayscale by averaging the three color channels, and the mean luminance was matched using the SHINE toolbox (Willenbockel et al., 2010).

The center scenes were always presented with a surrounding scene, except in the baseline block, where the central stimulus was shown alone (see below). Scene images were selected from two superordinate categories: indoor (restaurants and museums) and outdoor (parks and residential areas), each featuring two basic-level categories, with two exemplars per basic-level category. The center-surround stimuli depicted natural scenes with varied categorical relationships between the center and surround (Figure 1). Specifically, we devised four conditions: (1) identical exemplar condition, where the center and surround images were identical; (2) different exemplar condition, where the center and surround images belonged to the same basic-level category (e.g., museums) but were not identical; (3) different basic-level category condition, where the center and surround images belonged to the same superordinate category (e.g., indoor) but were from different basic-level categories; and (4) different superordinate category condition, where the center and surround images belonged to different superordinate categories.

**Figure 1:**
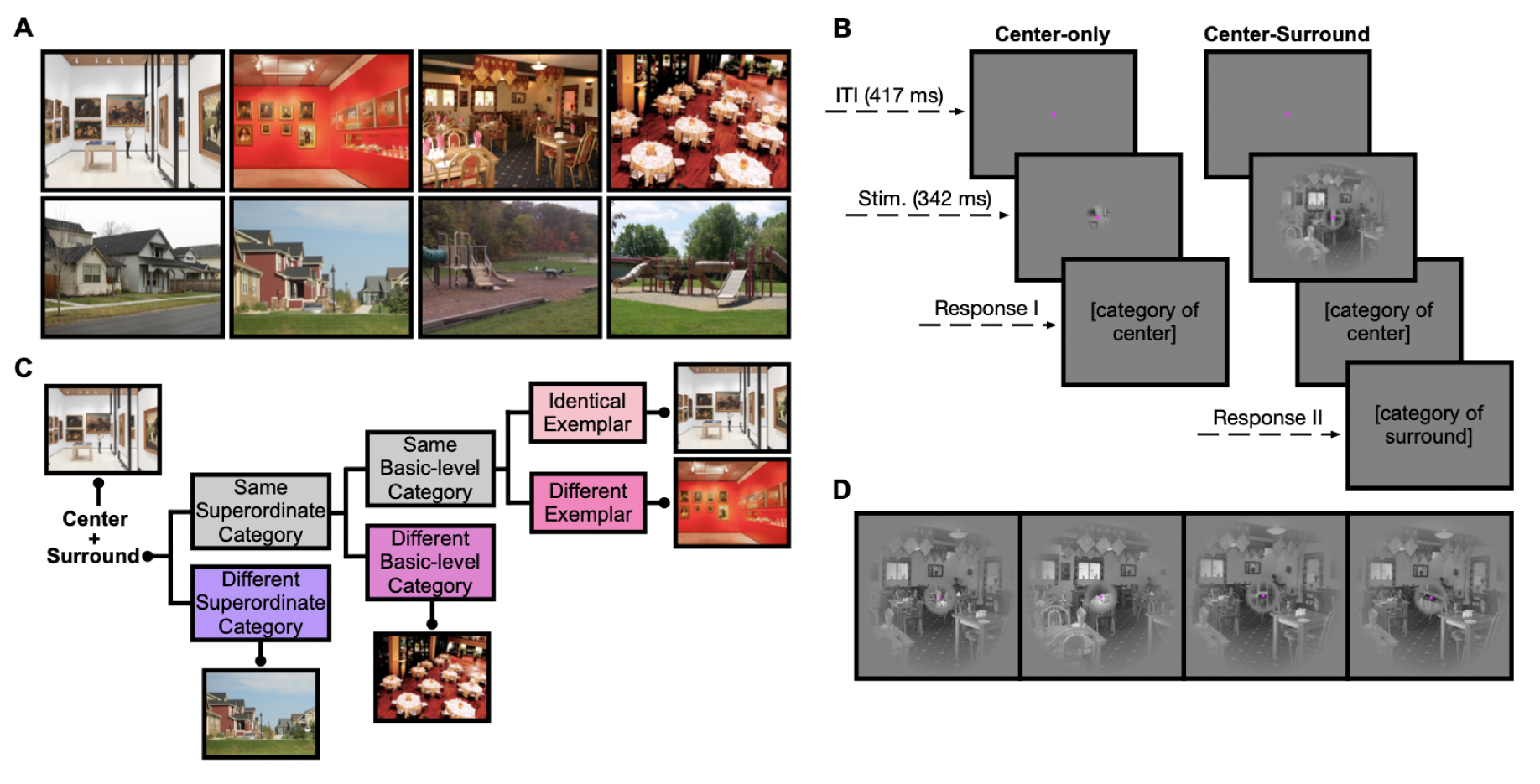
Stimuli and Paradigm. A) The natural scene images used to create the stimuli, drawn from two superordinate categories (indoor, outdoor) and two basic-level categories within each (restaurants, museums, parks, residential areas), with two exemplars per basic-level category. B) Experimental design for the Center-Only and Center+Surround blocks. Each trial began with a fixation point, followed by stimulus presentation, with the contrast level adjusted using 1-up 3-down staircase. In the Center-Only block, participants reported the superordinate category of the central scene. In the Center+Surround block, they first reported the category of the center scene, then the category of the surrounding scene. C) Center-surround stimuli were created for four conditions: (1) Identical Exemplar, where center and surround images were identical; (2) Different Exemplar, where images were from the same basic-level category but not identical; (3) Different Basic-Level Category, where images were from different basic-level categories within the same superordinate category; and (4) Different Superordinate Category, where images were from different superordinate categories (e.g., indoor vs. outdoor). D) Example stimuli for each category condition, shown from left to right: Identical Exemplar, Different Exemplar, Different Basic-Level Category, Different Superordinate Category.

The center-surround stimuli were presented in a single *Center-Surround* block. In half of the trials, the center and surround moved in the same direction; in the other half, they moved in opposite directions. The four category conditions (identical exemplar, different exemplar, different basic-level category, and different superordinate category) and two motion direction conditions (same-direction and opposite-direction) were presented in randomized order.

Additionally, a separate *Center-Only* block was included as a baseline, in which the central stimulus was presented alone without a surrounding scene. This baseline block allowed us to quantify the suppressive effect of adding the surround to the central stimulus.

### Procedure

Stimuli were presented using the Psychophysics toolbox (Brainard, 1997) with MATLAB R2022b (MathWorks, Natick, MA) on an AORUS FI32Q monitor (resolution 2560 × 1440, refresh rate 120 Hz). Participants were seated in a dark room with their heads stabilized using a chin rest at a distance of 60 cm from the monitor. The stimuli were presented foveally on a mid-gray background (128 RGB luminance). Responses were collected via a standard computer keyboard.

The experiment began with a practice session to allow participants to familiarize themselves with the task, followed by two main blocks: a Center-Only block and a Center+Surround block. The full session lasted approximately 100 minutes. Short breaks were provided during the experiment.

Participants were instructed to keep their eyes on a fixation point throughout the trial. On each trial, participants viewed foveally presented scene videos and performed a scene categorization task via keypress. Each trial began with a 416.6 ms fixation point followed by the presentation of the stimulus for 341.7 ms. In the Center-Only block, participants were asked to report the superordinate category of the central scene (i.e., indoor vs. outdoor). In the Center+Surround block, participants were instructed to attend to both the center and surrounding scenes. They first reported the category of the center scene, then reported the category of the surrounding scene. The second task was included to ensure that participants attended to both the center and the surround.

In all conditions, the Michelson contrast of the scenes was adjusted using an adaptive staircase procedure. In the Center-Only block, the contrast of the central scenes was adjusted using two interleaved 1-up, 3-down staircases, based on participants’ categorization judgments of the center scene. Importantly, two separate staircases were used: one starting from a very low contrast level (10%), making the task relatively difficult, and the other starting from a very high contrast level (90%), making the task relatively easy. Each staircase contained 160 trials, resulting in a total of 320 trials per participant for the baseline block.

In the Center+Surround block, the contrast of both the center and surrounding scenes was adjusted using independent 1-up, 3-down staircases, based on participants’ judgments of the center scene category. However, judgments about the surrounding scene category were not used for the contrast adjustment; they were used solely to direct attention to both center and surround. Importantly, the Center+Surround block always followed the Center-Only block, as the contrast threshold obtained from the baseline block was used as the starting point for the adaptive staircase procedure in the Center+Surround block. One adaptive staircase was used for each condition, yielding 8 staircases (4 category conditions × 2 motion directions). Each staircase included 160 trials, for a total of 1280 trials per participant in the Center+Surround block.

### Data Analysis

The contrast thresholds (corresponding to a 79% accuracy rate) were calculated for each participant and experimental condition by fitting a Gumbel psychometric function to the responses using the Palamedes toolbox (Kingdom & Prins, 2010) in MATLAB R2022b (MathWorks, Natick, MA). In cases where the psychometric fit failed to converge or yielded unreliable estimates (e.g., parameter values exceeded acceptable limits), thresholds were computed as the mean of the final six staircase reversals. This alternative method provides a robust approximation, as the reversal points tend to cluster around the observer’s threshold (Levitt, 1971).

Using these thresholds, we computed a suppression index (SI) to quantify the strength of surround suppression:

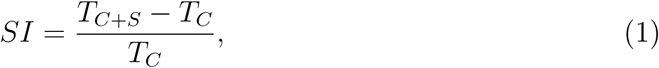

where *T_C_* and *T_C_*_+_*_S_* are the contrast thresholds for center-only and center+surround conditions, respectively. Positive SI values indicate suppression, negative values indicate facilitation, and an SI of zero indicates no contextual influence.

SI values exceeding ±2.5 standard deviations from the mean of the group were excluded from the analysis. Surround modulation was assessed by testing SI values against zero using two-tailed one-sample *t* -tests, with FDR correction applied for multiple comparisons.

To examine whether the suppression strength was influenced by the categorical similarity between center and surround scenes across different direction conditions, we ran linear mixed-effects models (LMMs), with SI values as the dependent variable. Categorical similarity (four levels: identical exemplar, different exemplar, different basic-level category, different superordinate category), motion-direction congruence (two levels: same-direction, opposite-direction), and their interaction were included as fixed effects, while between-subject variability was modeled using a random intercept.

All analyses were performed with custom MATLAB R2022b scripts (MathWorks, Natick, MA), except for the linear mixed-effects analysis, which was conducted in SPSS Statistics Version 25 (IBM Corp., Armonk, NY).

## Results

Figure 2 shows suppression indices (*SI*, Methods) derived from contrast thresholds. In Experiment 1, the presence and characteristics of the surround altered the perceived contrast of the central stimulus. A linear mixed-effects model revealed significant main effects of category (*F* (3,166)= 11.19, *p <* 0.001), and motion direction (*F* (1,166)= 9.33, *p* = 0.003). However, the interaction between category and direction was not significant (*F* (3,166)= 0.16, *p* = 0.92).

**Figure 2:**
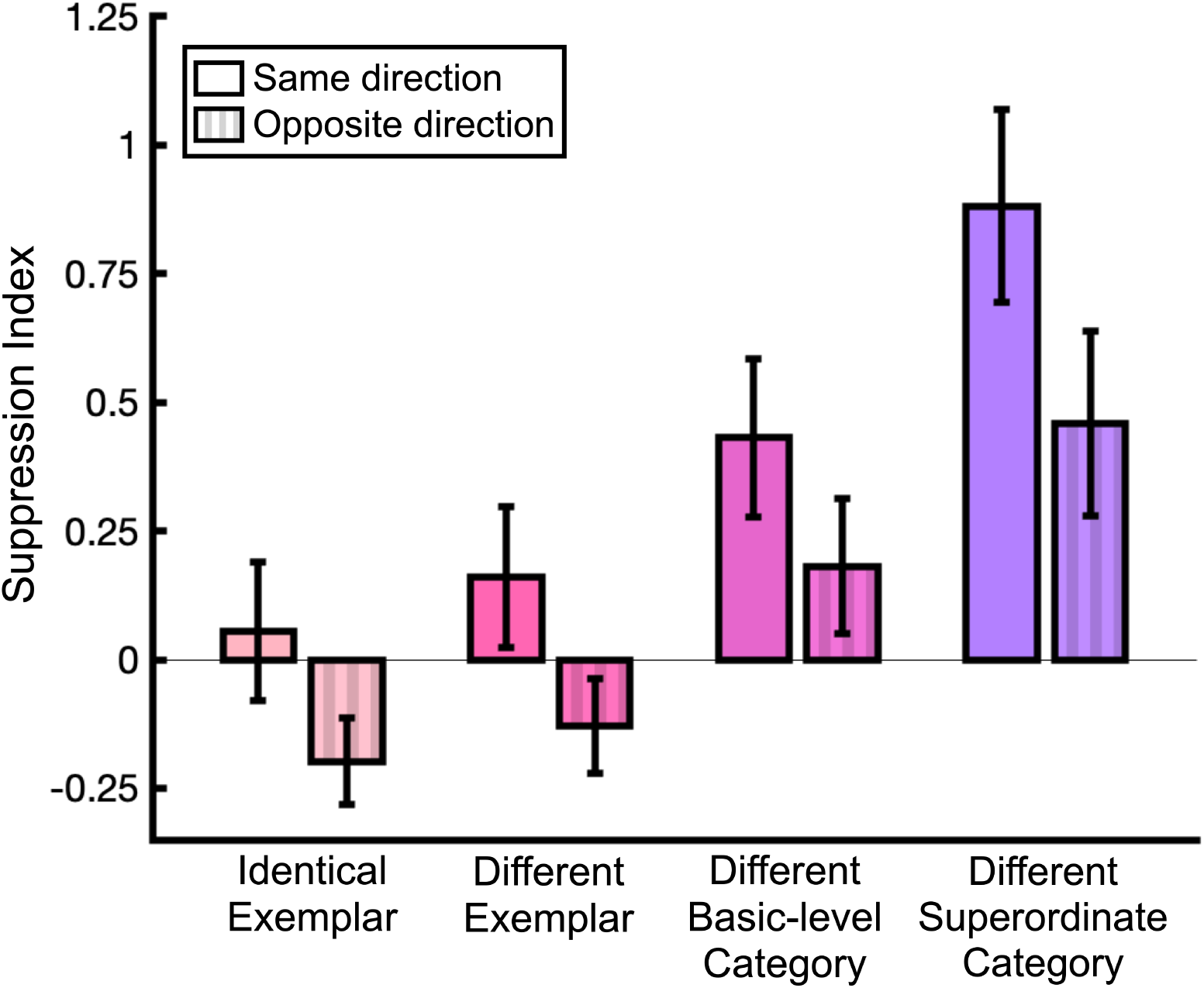
Suppression Index (SI) values from Experiment 1. SI values were computed for each participant across four category conditions (identical exemplar, different exemplar, different basic-level category, and different superordinate category) and two motion direction conditions (same- and opposite-direction). Higher SI values indicate stronger surround suppression, while negative SI values reflect surround facilitation. Error bars represent the standard error of the mean (SEM).

These results indicate that surround suppression was modulated by the motion direction of the surrounding stimuli. Consistent with previous findings (Kiniklioglu & Boyaci, 2022; Paffen, Alais, & Verstraten, 2005; Paffen, van der Smagt, et al., 2005), suppression was stronger when the center and surround moved in the same direction compared to opposite directions.

More importantly, across both motion conditions, suppression strength increased as categorical similarity between the center and surround decreased, with the strongest suppression observed when the center and surround belonged to different superordinate categories. Post hoc paired*−*sample *t* -tests with FDR correction confirmed that suppression was significantly greater in the different superordinate category condition than in all other category conditions (all *t* (21) *>* 2.69, *p*s *<* 0.02). Similarly, suppression was significantly stronger for different basic-level category condition than in the different exemplar and identical exemplar conditions (all *t* (21) *>* 2.80, *p*s *<* 0.02). In contrast, suppression strength did not differ between the identical and different exemplar conditions (*t* (21) = 1.34, *p* = 0.20).

Moreover, SI values were significantly greater than zero in the different superordinate category conditions for both same- and opposite-direction trials, as well as in the different basic-level category conditions for same-direction trials (*p*s *<* 0.05). These findings show that surround suppression is evident in dynamic natural scenes, particularly when the center and surround belong to different categories, and that suppression becomes stronger as categorical similarity decreases.

## Experiment 2

The results of Experiment 1 showed that the suppressive effect of the surround increased as the categorical similarity between the center and surround scenes decreased. One potential explanation for the relatively weak suppression in conditions with higher center-surround similarity is a possible order effect resulting from presenting the Center-Only block before the Center+Surround block. Although such an effect is unlikely to alter the overall pattern across conditions, it could bias the Suppression Index values toward facilitation. Additionally, presenting center-only and center+surround trials in separate blocks may have introduced confounds related to spatial attention. To address these concerns, Experiment 2 was designed to replicate Experiment 1 while eliminating potential confounding factors. To this end, we presented the center-only and center+surround trials together in a single mixed block with randomized trial order.

## Methods

### Participants

Twenty-five volunteers (7 male, mean age = 28.7 years) participated in Experiment 2. All participants provided written informed consent and received monetary compensation for their participation.

### Stimuli and Procedure

The apparatus, stimuli, and overall setup were identical to those used in Experiment

1. The key change in Experiment 2 was the inclusion of center-only trials within the Center+Surround block. Specifically, center-only and center+surround trials were randomly presented within a single Mixed block to control for potential order effects and attentional confounds introduced by separate block presentations.

The general procedure remained the same as in Experiment 1. The experiment began with a brief practice session, followed by a separate Center-Only block used to determine the starting values for the adaptive staircases. This was followed by the Mixed block. In the Mixed block, participants were instructed to report only the center scene category on center-only trials and to report the center category first, followed by the surround category, on center+surround trials. Participants were informed in advance that both trial types (center-only and center+surround trials) would appear in randomized order within the same block.

In the Mixed block, stimulus contrast for both center and surround images was adjusted using 1-up, 3-down staircases based on responses to the center scene category. One adaptive staircase was used per condition, resulting in a total of nine staircases (four category conditions × two motion directions plus one for the center-only trials). Each staircase started at the threshold value obtained from the initial Center-Only block.

### Data Analysis

As in Experiment 1, contrast thresholds corresponding to 79% accuracy were estimated using the Palamedes toolbox (Kingdom & Prins, 2010) in MATLAB R2022b (MathWorks, Natick, MA). Suppression Index (SI) values were calculated separately using two different baselines: (1) center-only trials from the Center-only block and (2) center-only trials from the Mixed block (see Experiment 1 Methods for details). SI values were computed for each participant and condition using both baselines. Further statistical tests were performed on the SI values.

We compared SI values against zero using one-sample, two-tailed Student’s *t* -tests, with FDR correction applied for multiple comparisons. Next, we analyzed the effects of categorical similarity and motion-direction congruence on suppression strength using linear mixed-effects models (LMMs), with SI values as the dependent variable. Categorical similarity, motion direction, and their interaction were included as fixed effects, while between-subject variability was modeled using a random intercept.

Additionally, to confirm that block structure did not influence baseline sensitivity, we compared contrast thresholds for center-only trials between the Center-only block and the Mixed block using a paired*−*sample *t* -test.

## Results

Figure 3 shows suppression indices (*SI*; Methods) calculated with center-only trials from the initial Center-only block. As in Experiment 1, the presence and the characteristics of the surround altered the perceived contrast of the central stimulus. Linear mixed-effects model (LMM) results showed a significant main effect of category (*F* (3,185)= 22.82, *p <* 0.001), and a significant main effect of motion direction (*F* (1,185)= 5.01, *p* = 0.03). However, the interaction between category and motion direction was not significant, *F* (3,185)= 1.97, *p* = 0.12).

**Figure 3:**
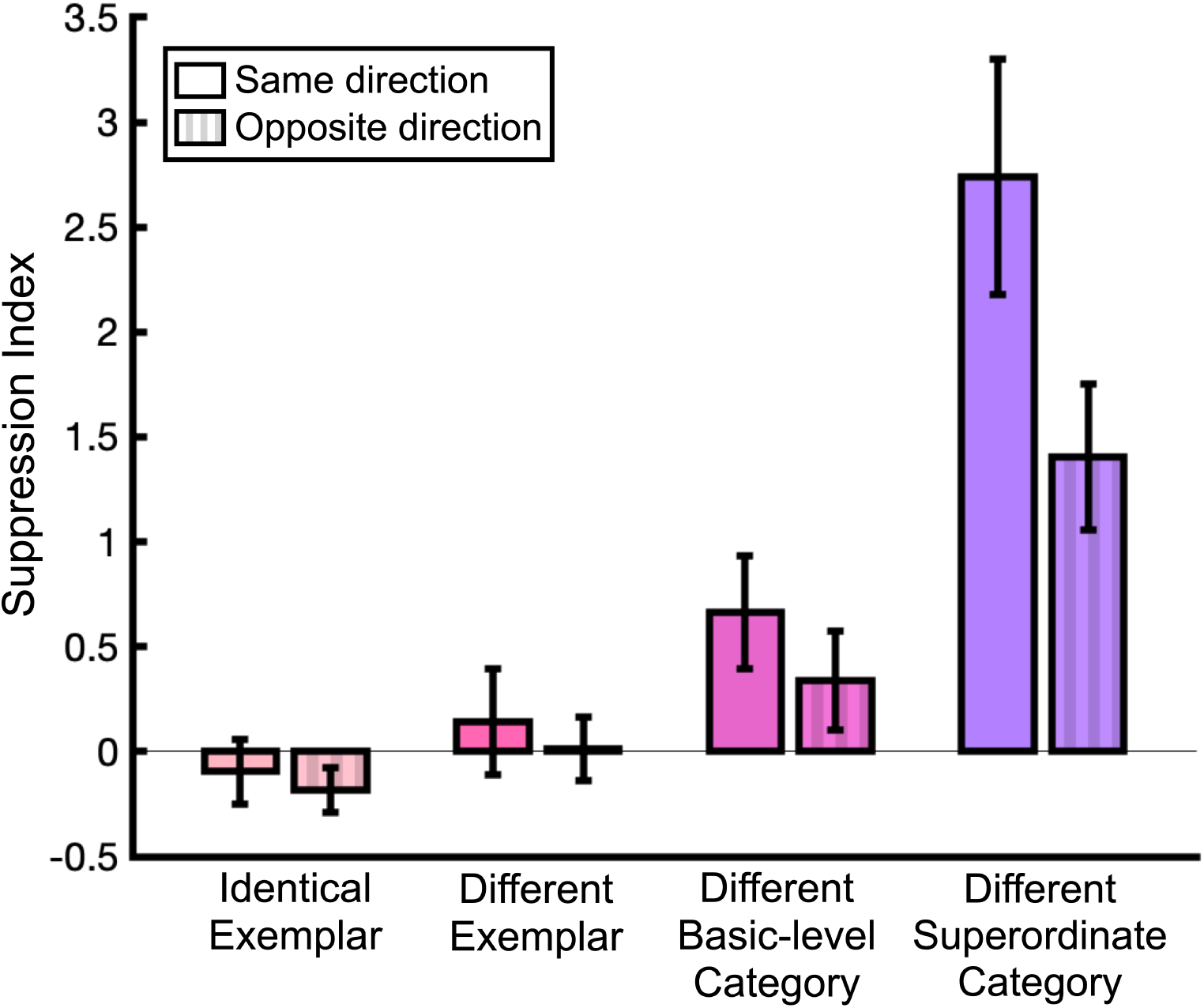
Suppression Index (SI) values from Experiment 2, calculated using centeronly trials from the Center-Only block. SI values were computed for each participant across four category conditions (identical exemplar, different exemplar, different basic-level category, and different superordinate category) and two motion direction conditions (same- and opposite-direction). Higher SI values indicate stronger surround suppression, while negative SI values reflect surround facilitation. Error bars represent the standard error of the mean (SEM).

As in Experiment 1, surround suppression was stronger when the center and surround moved in the same direction. Moreover, across both motion directions, SI values increased as categorical similarity between center and surround scenes decreased, replicating the pattern observed in Experiment 1. Post hoc paired*−*sample t-tests with FDR correction confirmed that suppression was significantly greater for the different superordinate category condition compared to other category conditions (all *t* (24) *>* 3.91, *p*s *<* 0.001). Likewise, suppression was significantly stronger in the different basic-level category condition than in the identical exemplar condition (*t* (24) = 4.32 all *p <* 0.001), and marginally stronger than in the different exemplar condition (*t* (24) = 2.06 all *p* = 0.06). In contrast, suppression strength did not differ between the identical and different exemplar conditions (*t* (24) = 1.45, *p* = 0.16). Moreover, SI values were significantly greater than zero in the different superordinate category conditions for both same- and opposite-direction trials, as well as in the different basic-level category conditions for same-direction trials (*p*s *<* 0.05).

To assess whether block order influenced baseline sensitivity, we compared contrast thresholds from center-only trials in the Center-Only and Mixed blocks. The threshold values were slightly higher in the Mixed block (*M* =0.19, *SD* =0.13) compared to the Center-Only block (*M* =0.18, *SD* =0.17). However, a paired*−*sample *t −*test showed no significant difference between blocks, (*t* (24)= *−*0.20, *p* = 0.84). Thresholds in the two blocks were moderately correlated across participants (*r* = 0.48, *p <* 0.05). These results suggest that baseline contrast sensitivity was relatively stable across blocks, although small differences might still influence the SI values.

Therefore, we recalculated SI values using center-only trials from the Mixed block (Figure 4) and re-ran the LMM to assess whether the pattern observed in Experiment 1 remained. Results again showed a significant main effect of category (*F* (3,185)= 26.29, *p <* 0.001) and a significant main effect of motion direction (*F* (1,185)= 4.74, *p* = 0.03). The interaction between category and motion direction was not significant, *F* (3,185)= 1.08, *p* = 0.36).

**Figure 4:**
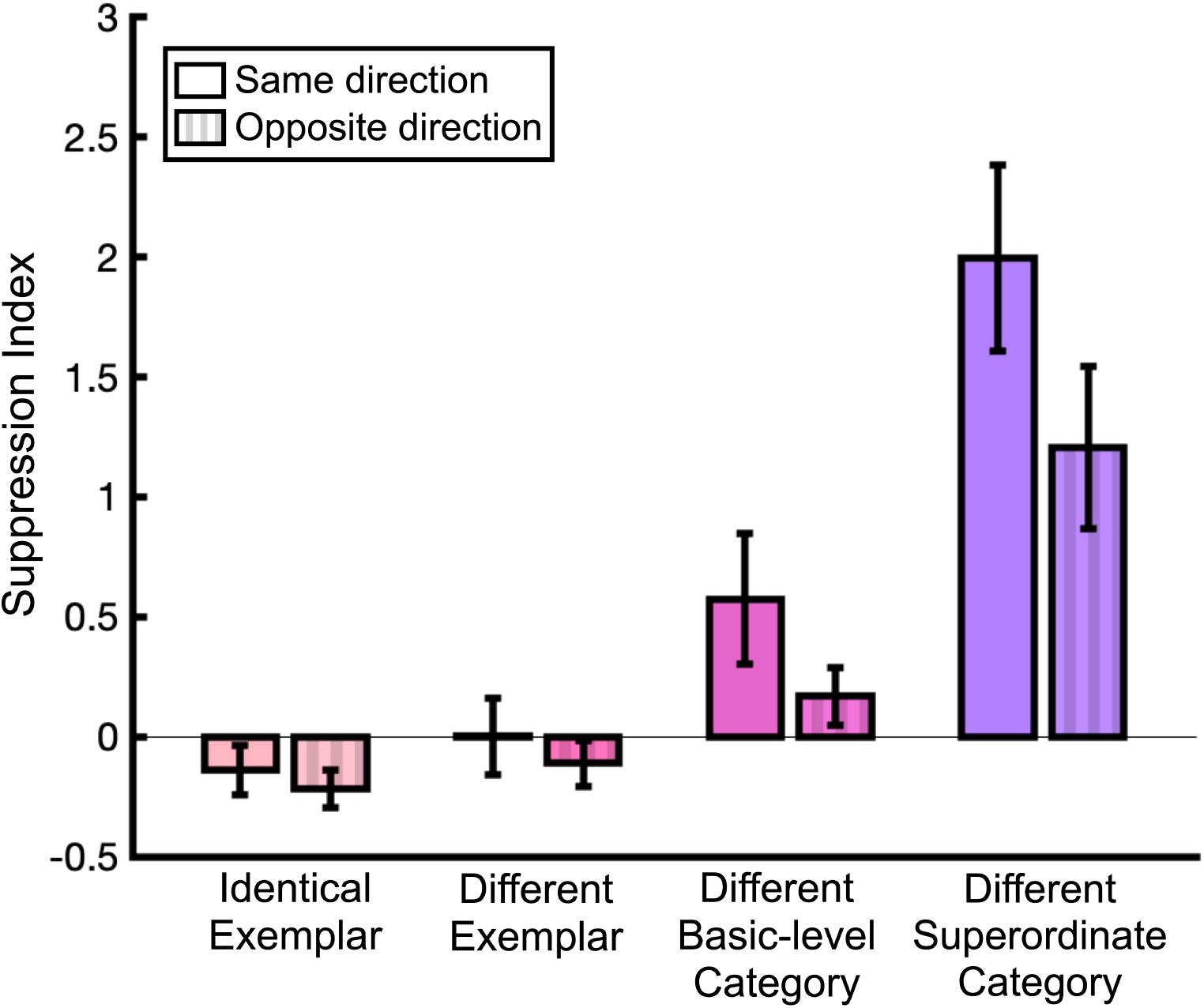
Suppression Index (SI) values from Experiment 2, calculated using center- only trials from Mixed block.SI values were computed for each participant across four category conditions (identical exemplar, different exemplar, different basic-level category, and different superordinate category) and two motion direction conditions (same- and opposite-direction). Higher SI values indicate stronger surround suppression, while negative SI values reflect surround facilitation. Error bars represent the standard error of the mean (SEM).

Taken together, these findings confirm that the effects of categorical similarity and motion congruence on surround suppression were robust across both experiments and were not driven by block structure or attentional confounds. The replication of the effects across block types suggests that the well-established influence of motion congruence on suppression, previously demonstrated with simple stimuli, also generalizes to complex naturalistic scenes. However, while physical similarity typically leads to stronger suppression with low-level stimuli (Cavanaugh et al., 2002a; Flevaris & Murray, 2015; Serrano-Pedraza et al., 2012; Shushruth et al., 2012), this was not the case for naturalistic stimuli. Instead, categorical dissimilarity between the center and surround produced stronger suppression, highlighting a key difference between suppression mechanisms for highly simplified versus naturalistic stimuli.

## Discussion

The current study investigated how motion congruence and categorical similarity between central and surrounding stimuli shape surround suppression during natural scene perception. Across two experiments, we measured contrast thresholds for categorizing foveally presented natural scenes while systematically varying the relationship between the center and surround across four levels of categorical similarity (identical exemplar, different exemplar, different basic-level category, different superordinate category) and two levels of motion congruence (same vs. opposite direction). We found that contrast sensitivity for the central scene was significantly modulated by the presence and characteristics of the surrounding scene. Surround suppression was strongest when the surround moved in the same direction as the center, replicating classic motion congruence effects found with simple stimuli (Allman et al., 1985; Cavanaugh et al., 2002b; Kastner et al., 1995; Kiniklioglu & Boyaci, 2025; Kiniklioglu & Boyaci, 2022; Lamme, 1995; Paffen, Alais, & Verstraten, 2005; Paffen, van der Smagt, et al., 2005). Importantly, we also found that suppression increased as the center and surround became more categorically dissimilar, with the strongest suppression occurring when they belonged to different superordinate categories (e.g., indoor vs. outdoor).

These results provide the first behavioral evidence of surround suppression in human perception using dynamic natural scenes. Although suppression effects have been widely documented with simple stimuli such as gratings (Cavanaugh et al., 2002b; Schallmo et al., 2018; Tadin et al., 2003), their role in processing complex ecologically valid visual stimuli in humans has remained unclear. A few animal studies using naturalistic stimuli have reported mixed results (Coen-Cagli et al., 2015; Fu et al., 2024; Onat et al., 2013). Our results show that surrounding scene context systematically reduces sensitivity to a central scene, establishing that surround suppression also occurs with naturalistic stimuli in human scene perception.

This suppression effect gets weaker, or may even disappear entirely, when the center and surround move in opposite directions. This has been shown in both neurophysiology studies (Allman et al., 1985; Born & Tootell, 1992; Cavanaugh et al., 2002b; Kastner et al., 1995; Lamme, 1995) as well as behavioral studies (Kiniklioglu & Boyaci, 2022; Paffen, Alais, & Verstraten, 2005; Paffen, van der Smagt, et al., 2005). Consistent with these findings, we observed stronger suppression when the center and surround moved in the same direction compared to when they moved in opposite directions. This effect was robust across all category conditions, suggesting that motion congruence modulates suppression even with complex, naturalistic stimuli. These results suggest that motion-based center-surround interactions, thought to reflect inhibitory mechanisms in the middle temporal cortex (MT hypothesis; see Tadin et al., 2003), generalize to more ecologically valid conditions.

In both experiments, suppression strength increased as the categorical distance between center and surround increased. Surrounds that belonged to different superordinate categories (e.g., indoor vs. outdoor) produced the strongest suppression, whereas scenes from the same basic-level category (e.g., two museums) produced weaker or no suppression. These results suggest that categorical similarity plays a critical role in contextual modulation: the more categorically dissimilar the surround, the more it interferes with the perception of the central scene. This pattern contrasts with findings from studies using low-level stimuli, where suppression typically increases with center-surround similarity in features like orientation or spatial frequency (Cavanaugh et al., 2002a; DeAngelis et al., 1994; Flevaris & Murray, 2015; Schallmo et al., 2016; Serrano-Pedraza et al., 2012; Shushruth et al., 2012; Walker et al., 1999). For example, the strongest suppression is observed when the center and surround are physically identical. In our study, however, no suppression was observed when the center and surround scenes were drawn from the same exemplar; instead, suppression increased with the greater categorical dissimilarity. These findings align with animal studies using natural stimuli, which show that perceptually similar surround stimuli facilitate neural responses in the cat visual cortex, whereas dissimilar surrounds lead to suppression (Fu et al., 2024; Onat et al., 2013). Taken together, our results suggest that, in natural vision, suppression is not solely driven by low-level feature similarity, but is also influenced by higher-level categorical structure.

Our findings also align with previous research showing that categorical incongruence between center and surround scenes impairs scene categorization accuracy (Faurite et al., 2024; Lukavský, 2019; Trouilloud et al., 2022). Critically, we extend these findings by demonstrating that such categorical congruency effects also modulate low-level perceptual sensitivity, as reflected in contrast thresholds, even when the task involves high-level scene categorization. Moreover, Peyrin et al. (2021) showed that even when central and peripheral scenes are categorically congruent (e.g., two man-made scenes), physical dissimilarities between the scenes (e.g., buildings vs. streets) lead to decreased central scene categorization accuracy. Consistent with this, we found suppression effects even when the center and surround belonged to the same task-relevant superordinate category (e.g., two indoor scenes) but differed at the basic level (e.g., restaurant vs. museum).

Furthermore, there was no interaction between motion congruence and categorical similarity. This suggests that these two types of contextual information modulate suppression independently. One possibility is that suppression arises from multiple stages of processing, with earlier visual areas primarily sensitive to basic features like contrast and motion (Angelucci et al., 2017), while higher-level perceptual systems contribute additional modulation based on categorical information. Together, our results suggest that traditional models of surround suppression, which have focused primarily on low-level features such as contrast, orientation, and motion (Angelucci et al., 2017), should be expanded to incorporate the influence of higher-level contextual factors.

In real-world environments, scenes generally follow consistent categorical structures (e.g., a kitchen containing related objects such as appliances, utensils, and cabinets, all organized within a coherent spatial layout). To process such environments efficiently, the visual system integrates spatial and semantic context across multiple levels (Bar, 2004; Kaiser et al., 2019; Vo, 2021). Some theoretical frameworks propose that the brain actively suppresses contextually incongruent input to enhance perceptual efficiency (Angelucci & Bressloff, 2006; Gilbert & Li, 2013; Rao & Ballard, 1999). In line with this, multiple studies have shown that coherent scene structure enhances or speeds up scene and object processing (Chen et al., 2022; Davenport & Potter, 2004; Kaiser & Peelen, 2018; Kaiser et al., 2019, 2020, 2021; Võ & Wolfe, 2013). Therefore, suppression mechanisms in natural contexts may help to prioritize coherent, meaningful input while filtering out conflicting or irrelevant information.

Our findings support this idea by demonstrating that incongruent surrounds (both at the basic-level and superordinate level) lead to suppression, whereas congruent surrounds do not. Although previous studies with simple stimuli have reported suppression with identical surrounds, typically explained as eliminating statistical redundancies (Angelucci et al., 2017; Coen-Cagli et al., 2012; Nurminen & Angelucci, 2014; Schwartz & Simoncelli, 2001; Vinje & Gallant, 2000), our results suggest that at higher levels of processing, redundancy may take a different form. In natural scenes, categorically incongruent information may be treated as irrelevant and suppressed, whereas with simple stimuli, identical information may be suppressed as redundant. Both mechanisms likely serve to prioritize task-relevant input by filtering out irrelevant or predictable contextual information. Thus, context-dependent suppression may contribute to perceptual stability by minimizing the impact of incongruent elements.

A common concern in studies of contextual modulation is the potential influence of spatial attention. To address this, we employed both blocked (Experiment 1) and mixed (Experiment 2) designs to control for possible order effects and attentional biases. The consistency of results across both designs suggests that the observed suppression effects cannot be attributed to block order or attentional strategies. One might also argue that the suppression effects observed in natural scenes reflect the increased cognitive demands of the second task. However, if task difficulty were the primary factor, suppression strength should not differ between same- and opposite-direction trials. Instead, our results consistently show stronger suppression under the same-direction condition.

Together, these findings support the interpretation that suppression in our paradigm reflects active, stimulus-driven contextual inhibition, rather than passive attentional withdrawal or task-related artifacts.

Although the present study provides robust evidence for both categorical and motion-based surround suppression in natural vision, several questions remain open for future research. First, we did not explicitly manipulate low-level visual features such as spatial frequency or amplitude spectrum, which may covary with categorical similarity and potentially contribute to suppression effects (Peyrin et al., 2021). Future studies could independently vary such physical properties and categorical coherence to better disentangle their influences. Second, although we observed reliable effects of categorical similarity, the underlying neural mechanisms remain to be clarified. Follow-up studies using neuroimaging methods could help localize the sources and time course of suppression and determine whether distinct neural circuits mediate motion-based and category-based suppression. Finally, in our experiments, participants performed a categorization task on a relatively narrow range of scene categories. It remains an open question whether similar suppression effects would generalize to a broader set of scenes, such as comparisons between natural and man-made environments, and beyond categorization tasks. Together, these directions highlight the need for further research to refine our understanding of how high-level contextual information shapes visual processing in natural vision.

## Conclusion

To conclude, our study demonstrates that surround suppression in natural scene perception is shaped by both motion-direction congruence and categorical similarity.

Consistent with results with simple stimuli, suppression was stronger when the center and surround moved in the same direction. However, unlike low-level studies, where identical center-surround features typically yield the greatest suppression, we observed no suppression in the identical exemplar condition. Instead, suppression increased as the categorical similarity between center and surround decreased. Thus, in natural vision, contextual modulation is driven not only by low-level feature similarity but also by higher-level categorical structure. These findings advance our understanding of contextual modulation in natural vision, and suggest that context-dependent suppression may help the visual system prioritize coherent, task-relevant information while suppressing incongruent input.

## Data Availability

The data and analysis code will be available on the Open Science Framework.

## Acknowledgements

This work was supported by the Deutsche Forschungsgemeinschaft (DFG), grants SFB/TRR 135 (project no. 222641018), KA4683/5-1 (project no. 518483074), KA4683/6-1 (project no. 536053998), and under Germany’s Excellence Strategy (EXC 3066/1 “The Adaptive Mind”, project no. 533717223). It was further supported by an European Research Council (ERC) Starting Grant (PEP, ERC-2022-STG 101076057). Views and opinions expressed are those of the authors only and do not necessarily reflect those of the European Union or the European Research Council. Neither the European Union nor the granting authority can be held responsible for them. The authors thank Tugce Dalmis for supporting the data collection for both Experiments. The authors declare no competing interests.

